# The fast diffusion of NKCC1 along the axon is driven by glutamatergic activity

**DOI:** 10.1101/2022.12.11.519960

**Authors:** Etienne Côme, Erwan Pol, Zaha Merlaud, Juliette Gouhier, Marion Russeau, Xavier Marques, Sophie Scotto-Lomassese, Imane Moutkine, Sabine Lévi

## Abstract

NKCC1 and KCC2 transporters regulate neuronal chloride homeostasis and thus synaptic inhibition. KCC2 activity is tuned by diffusion-capture in an activity-dependent manner. The mechanisms controlling NKCC1 in neurons are unknown. We found using super-resolution imaging that NKCC1 like KCC2 form nanodomains in the somato-dendritic membrane at extrasynaptic sites and at the periphery of excitatory and inhibitory synapses. NKCC1 nanoclusters are half the size and density in molecules than KCC2 clusters. This is accompanied by a higher mobility of NKCC1 compared to KCC2 in the dendritic membrane, suggesting a weaker NKCC1 anchoring to the cytoskeleton. In contrast, NKCC1, but not KCC2, is confined to endocytic zones, which would explain its controlled surface expression and the fact that endocytic zones would provide a reservoir from which NKCC1 could be released into the membrane. Finally, we show an increased confinement of NKCC1 in axons but not in dendrites upon glutamatergic activity blockade, indicating a selective mechanism of regulation in the axon. We propose that a rapid regulation of NKCC1 by lateral diffusion in the axon would control presynaptic glutamate release and the firing of action potentials.

## Introduction

Synaptic inhibition in the somato-dendritic compartment is mediated by chloride permeable GABA type A receptors (GABA_A_Rs) in the mature brain, whereas in the early stages of development, GABA_A_R in this compartment is depolarizing/excitatory (ref in Kaila et al., 2014). The shift from depolarizing to hyperpolarizing GABA_A_R-mediated responses during the second postnatal week in rodents is mainly due to the expression of the chloride exporter KCC2 (Rivera et al., 1999). The fact that NKCC1 is expressed at a constant level throughout life in different cell types and subdomains (ref in Virtanen et al., 2020) raises the question of its role in the mature brain. The contribution of NKCC1 to chloride homeostasis in mature neurons seems negligible, at least under physiological conditions. The presence of NKCC1 in the axon initial segment (AIS) of mature pyramidal cells at axo-axonal GABAergic synapses formed by chandelier interneurons maintains a depolarized E_GABA_ value (Khirug et al., 2008). By exciting presynaptic GABA_A_R, GABA release from interneurons would facilitate glutamate release from presynaptic terminals of pyramidal cells (Jang et al., 2006). NKCC1 is also expressed by non-neuronal cells such as astrocytes (Henneberger et al. 2020; Wilson and Mongin, 2019). In glioblastoma, the interaction of NKCC1 with cofilin by regulating actin dynamics would provide a mechanism for cell migration (Schiapparelli et al. 2017). In the context of LTP, the interaction of NKCC1 with cofilin would regulate the dynamics of astrocytic processes via combined actions on the actin network and water influx, which would allow glutamate spillover and LTP (Henneberger et al. 2020). Therefore, the main contribution of NKCC1 in the mature brain is to regulate the presynaptic release of neurotransmitter and synaptic plasticity through regulation of astrocytic shape.

Key cellular and molecular mechanisms regulate the membrane turnover of KCC2. They involve activity-dependent regulation of the transporter membrane turnover and endocytosis (Côme et al., 2019a, 2019b). Our team has shown the contribution of lateral diffusion in the rapid control of KCC2 membrane stability and Cl^-^ neuronal homeostasis in response to changes in neuronal glutamatergic and GABAergic activities (Chamma et al., 2013; Heubl et al., 2017). The transporter alternates between periods of confinement within clusters near synapses and periods of free movement outside the clusters. The free transporter can then be targeted to endocytic zones where it is internalized and then degraded or recycled to the plasma membrane. We proposed that these different pools of transporters are in dynamic equilibrium, allowing the fine-tuning of synapses in response to local fluctuations in synaptic glutamatergic and GABAergic activities (Côme et al., 2019a, 2019b). Since changes in KCC2 mobility occur within tens of seconds (Heubl et al., 2017), lateral diffusion is probably the first mechanism modulating KCC2 stability and function in the membrane.

Unlike KCC2, almost nothing is known about the cellular and molecular mechanisms regulating NKCC1 in neurons. Here we compared KCC2 and NKCC1 membrane organization and we studied the contribution of the diffusion-capture mechanism in the regulation of NKCC1 at the surface of axons and dendrites in mature hippocampal neurons.

## Results

### NKCC1a and NKCC1b are targeted to somato-dendritic and axonal compartments of mature hippocampal neurons

Two splicing variants of NKCC1 have been described: NKCC1a and NKCC1b (Randall et al., 1997). NKCC1b differs from NKCC1a by the absence of an exon that contains a putative consensus sequence for PKA phosphorylation (Randall et al., 1997) and a dileucine motif essential for the basolateral sorting of NKCC1a in polarized epithelial cells (Carmosino et al., 2008). In the absence of antibodies labeling NKCC1a and NKCC1b, we expressed in primary cultures of hippocampal neurons, NKCC1a or NKCC1b constructs bearing a 2xHA tag at position Histidine^398^ in the second extracellular loop, a 3xFlag tag and a monomeric Venus tag inserted at the N-terminus (Somasekharan et al., 2013). We report using surface HA labeling and confocal microscopy in mature hippocampal neurons at 21 days in vitro (DIV) that NKCC1a (Fig. 1 A-B) and NKCC1b (Fig. 1 C-D) are targeted to the somato-dendritic plasma membranes. This somato-dendritic localization of NKCC1a and NKCC1b is similar to that of KCC2 (Fig. 1 E-F). The fact that NKCC1a and NKCC1b carrying or not the dileucine motif are both detected in the neuronal cell body and along dendrites (Fig. 1 A and C, respectively) indicates that the dileucine motif is not essential for the somato-dendritic localization of the transporter. Labeling of NKCC1a, NKCC1b, and KCC2 was performed in parallel allowing to compare them. We observed that the labeling of NKCC1a (Fig. 1 A-B) and NKCC1b (Fig. 1 C-D) is weaker than that of KCC2 (Fig. 1 E-F) on the surface of mature hippocampal neurons. This means that **mature neurons express less NKCC1a and NKCC1b on their surface than KCC2.**

**Fig. 1.**
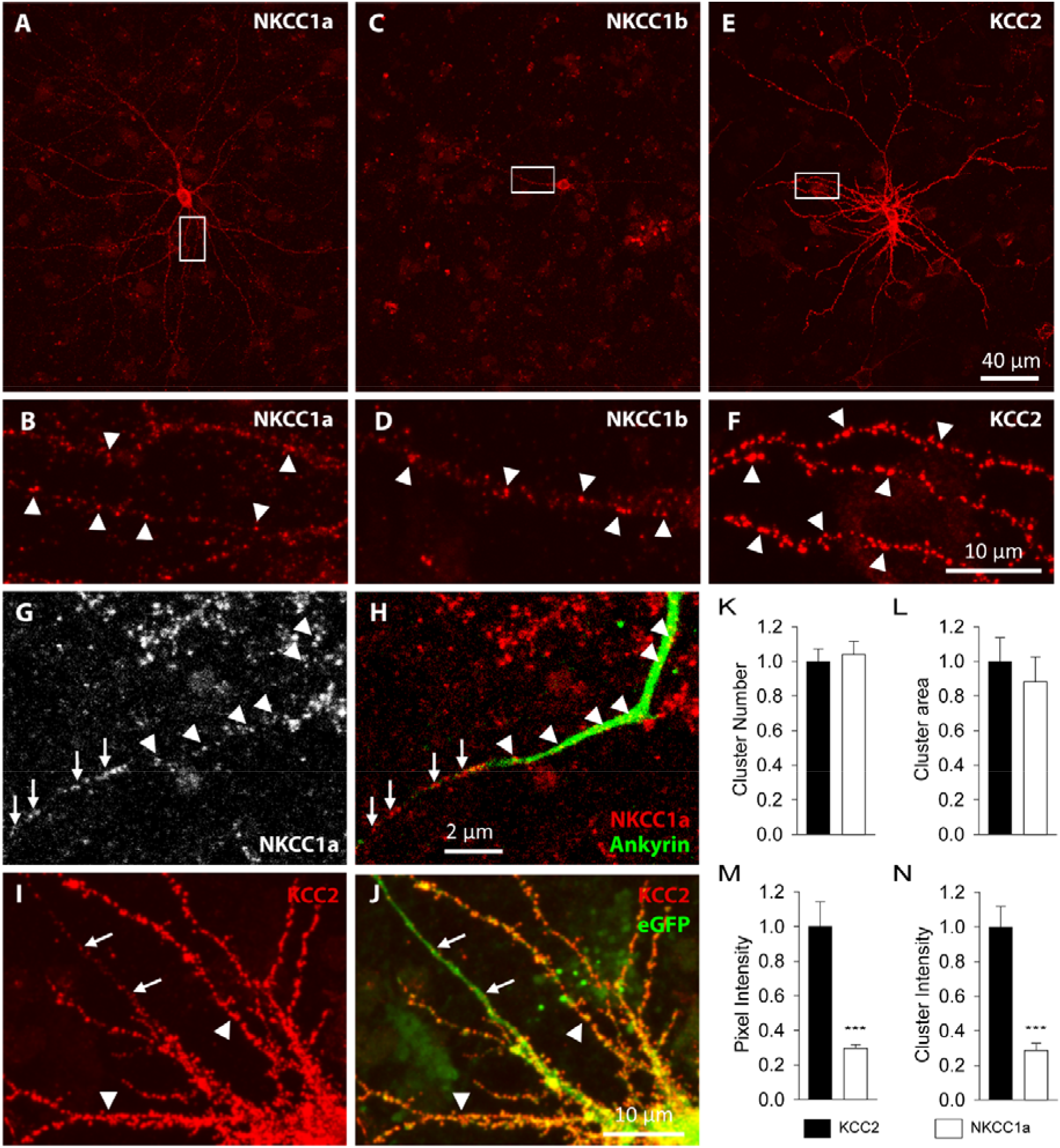
NKCC1a and NKCC1b are clustered in the somato-dendritic and axonal compartments of mature hippocampal neurons. **A-F**, Surface staining of NKCC1a (A-B), NKCC1b (C-D) and KCC2 (E-F) in 21 DIV old hippocampal neurons transfected with NKCC1a-HA, NKCC1b-HA or KCC2-Flag constructs. Scale bars: A, C, E, 40 μm; B, D, F, 10 μm. Note that the labelling of NKCC1a and NKCC1b is weaker than that of KCC2 at the surface of the soma and of neurites of mature hippocampal neurons. B, D and F correspond to the magnification of the dendritic regions outlined on the neurons in A, C and E, respectively. Note that NKCC1a, NKCC1b and KCC2 form many clusters along the dendrites (arrowheads) and that the intensity of KCC2 clusters is stronger than that of NKCC1a and NKCC1b clusters. **G-H**, Double-labelling of NKCC1a (white in G, red in H) and ankyrin (green in H) in mature neurons. Scale bar: 2 μm. NKCC1a clusters are detected in the initial segment of the axon (arrowheads) identified by ankyrin immunoreactivity and along the axon remote from the AIS (arrows). **I-J**, Immunodetection of membrane-associated KCC2 with anti-Flag antibody **(I)**in a neuron transfected with KCC2-Flag and cytoplasmic eGFP **(J)**. Scale bar: 10 μm. Note that KCC2 clusters are detected along the dendrites (arrowheads) of the transfected neuron but KCC2 labelling is absent from the thin GFP-positive neurite identified as the axon (arrows). **K-N**, Quantifications of NKCC1a (white bars) and KCC2 (black bars) cluster number (**K**), cluster area (**L**), intensity per pixel in the cluster (**M**), and intensity per cluster (**N**) showing reduced pixel and cluster intensity of NKCC1a clusters compared to KCC2. Data shown as mean ± SEM. In all graphs, values were normalized to the corresponding KCC2 mean values. KCC2 n = 22 cells, NKCC1a, n = 17 cells. Cluster Nb p = 0.62, area p = 0.29, pixel intensity p < 1.0 10^-3^, cluster intensity p < 1.0 10^-3^, 2 cultures. ***, p<1.0 10^-3^ (Mann–Whitney rank sum test).

We next assessed the localization of NKCC1a in the axon initial segment (AIS) and along the axon in double-staining experiments with ankyrin, a marker of the AIS (Fig. 1 G-H). As expected, ankyrin accumulates in the first ~10-100 μm of the axon (Fig. 1H). NKCC1a immunoreactivity was detected in the AIS (arrowheads in Fig. 1 G-H). However, NKCC1a was also present all along the axon (arrows in Fig. 1 G-H), indicating that its expression is not restricted to the AIS. This axonal localization of NKCC1a differs from that of KCC2, which is detected in the somato-dendritic compartment and excluded from the axon (Fig. 1 I-J), as shown before (Chamma et al., 2013). **The fact that NKCC1a and NKCC1b but not KCC2 are targeted to the axon suggests functional differences. This is in agreement with the E_GABA_ being depolarized at the AIS due to NKCC1 expression (Khirug et al., 2008) and KCC2 exclusion from the axon (Hübner et al., 2001; Williams et al., 1999).**

### Comparison of NKCC1 and KCC2 clustering in the dendrite

We next studied the subcellular distribution of NKCC1a and NKCC1b in mature hippocampal neurons. We found that NKCC1a and NKCC1b are not homogeneously distributed at the surface of the neurites of 21 DIV-old neurons but form clusters in the somato-dendritic membranes (Fig. 1 B and D, respectively). NKCC1 clusters are also found at the surface of the AIS and along the axon (Fig. 1 G-H). KCC2 also form clusters at the surface of the dendrites of hippocampal neurons (Fig. 1 F, I-J), as shown before (Chamma et al., 2013; Heubl et al., 2017; Watanabe et al., 2009).

We then quantified NKCC1a and KCC2 clustering in the dendrite. We found a similar density of NKCC1a and KCC2 clusters at the surface of neurons (Fig. 1K). NKCC1a and KCC2 clusters were of similar size (Fig. 1L). However, the mean pixel intensity per cluster (Fig. 1M) as well as the mean intensity per cluster (Fig. 1N) were significantly lower for NKCC1a, as compared to KCC2. Since NKCC1a and KCC2 cluster size are at the limit of the resolution of a standard epifluorescence microscope, we further compared NKCC1a and KCC2 clustering using super-resolution STORM. NKCC1a (Fig. 2 A-C) and KCC2 (Fig. 2 D-F) clusters were evenly distributed along the dendritic shaft (Fig. 2 A-B and D-E, respectively) and in dendritic spines (Fig. 2 A-C and D-F, respectively). Most NKCC1a clusters were round (arrowheads in Fig. 2A) while KCC2 clusters displayed round (arrowheads in Fig. 2D) or elongated shapes (arrows in Fig. 2D). We quantified these observations and found that NKCC1a clusters were ~2-fold smaller than KCC2 clusters. The mean size of NKCC1a clusters was 51.1 ± 1.6 nm^2^ (mean ± SEM) while KCC2 clusters had an average surface of 100.2 ± 3.5 nm^2^ (Fig. 2G). These NKCC1a nanoclusters were also half as dense in molecules as KCC2 clusters. On average, 111.1 ± 18.9 (mean ± SEM) unique molecules were detected per NKCC1a nanocluster in our experimental conditions whereas this number was 213.4 ± 7.4 for KCC2 (Fig. 2H). The detection density, i.e., the number of molecules detected per square micrometer, was also significantly lower for NKCC1a than for KCC2 (Fig. 2I), highlighting a lower compaction of NKCC1a clusters. **Therefore, we propose that NKCC1a form nanoclusters that recruit fewer molecules than KCC2 clusters.**

**Fig. 2.**
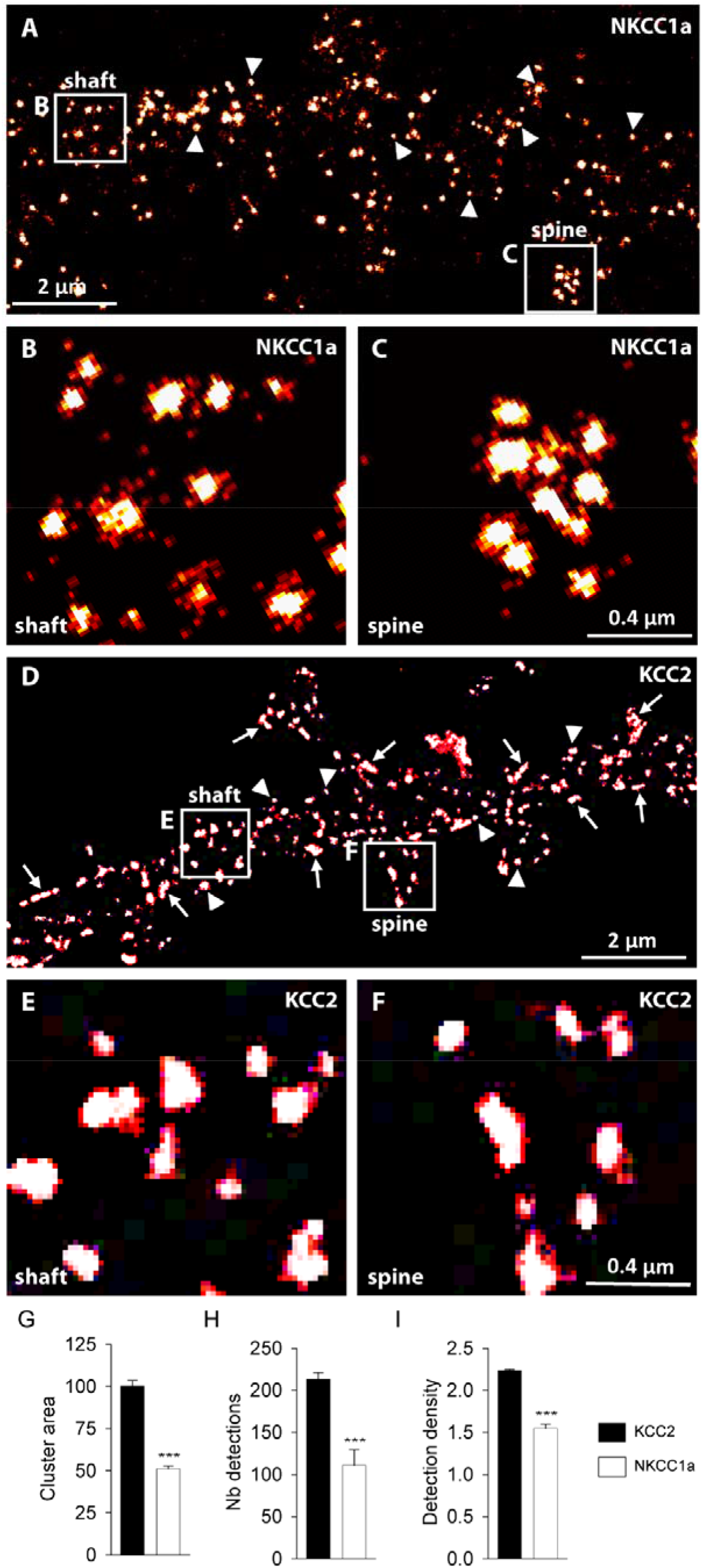
Nanoscale organization of NKCC1a and KCC2 transporters at the dendritic surface. **A-F**, STORM image reconstruction of membrane-associated NKCC1a (A-C) and KCC2 (D-F) in hippocampal neurons at 21 DIV. B-C, and E-F correspond to the magnification of the shaft and spines outlined on the neurons in A, D respectively. Scale bars: 2 μm in A, D; 0.4 μm in B-C and E-F. Note that NKCC1a and KCC2 form many nanoclusters of round shape (arrowheads) along the dendritic shaft and in spines. Some elongated (arrows) clusters of KCC2 are observed. **G-I**, Quantifications of NKCC1a (white bars) and KCC2 (black bars) cluster area (**G**), number of single molecules per nanocluster **(H)**and density of molecules per nanocluster **(I)**show reduced clustering for NKCC1a as compared to KCC2. Data shown as mean ± SEM. In all graphs, values were normalized to the corresponding KCC2 mean values. KCC2 n = 7491 nanoclusters, NKCC1a n = 1315 nanoclusters, D p < 1.0 10^-3^, E p < 1.0 10^-3^, F p < 1.0 10^-3^, 2 cultures. ***, p<1.0 10^-3^ (Mann–Whitney rank sum test).

### NKCC1a clusters are localized at the periphery of inhibitory synapses

To study the precise relationship of NKCC1a with inhibitory synapses, we performed STORM of NKCC1a HA labeling and PALM analysis of dendra2-gephyrin, a postsynaptic marker of inhibitory synapses. Dendra2-gephyrin detected inhibitory subsynaptic domains (SSDs) in dendritic shafts and in some spines of mature hippocampal neurons (Fig. 3 A-E). This is reminiscent of the nanoscopic organization of inhibitory postsynaptic densities shown using super-resolution (Battaglia et al., 2018; Crosby et al., 2019). NKCC1a nanoclusters were present in dendritic shafts and spines (Fig. 3 A). Many of these nanoclusters were at distance of dendra2-gephyrin SSDs (Fig. 3 A), indicating that they are in the extrasynaptic membrane and/or at excitatory glutamatergic synapses. However, NKCC1a clusters could also be detected near gephyrin SSDs (Fig. 3 A-C). Interestingly, NKCC1a nanoclusters do not colocalize with gephyrin SSDs but are distributed around gephyrin SSDs (Fig. 3 A-C). This indicates a perisynaptic rather than postsynaptic localization of NKCC1a. This distribution of NKCC1a is similar to that of KCC2. KCC2-Flag STORM and dendra2-gephyrin PALM reveal the presence of KCC2 clusters in dendritic shaft and spines at distance of inhibitory synapses (Fig. 2D) and a perisynaptic localization of KCC2 aggregates at inhibitory synapses (Fig. 2 D-E). **This perisynaptic accumulation of chloride transporters at inhibitory synapses would allow rapid adjustment of neuronal chloride homeostasis in response to changes in GABAergic transmission.**

**Fig. 3.**
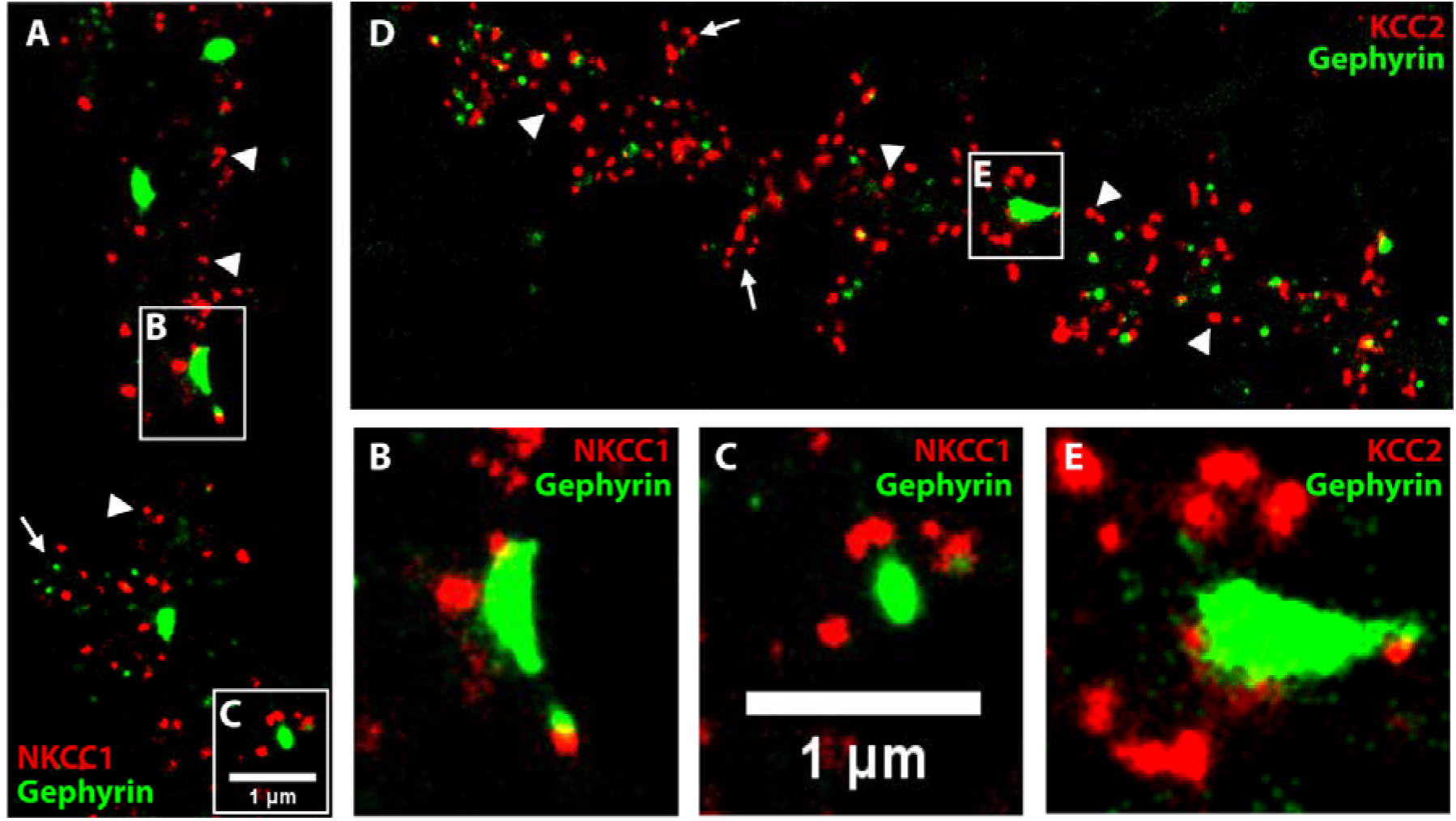
NKCC1a and KCC2 nanoclusters are surrounding inhibitory synapses. **A-E**, 2-color STORM of NKCC1a (red in A-C) or KCC2 (red in D-E) and PALM of gephyrin (green in A-E) in 21 DIV-old neurons. B-C and E correspond to the magnification of the synapses outlined on the neurons in A and D, respectively. Scale bars: 1 μm. Note that many NKCC1a and KCC2 nanoclusters surround inhibitory synapses formed on the dendritic shaft. Numerous NKCC1a nanoclusters are also detected at distance from the inhibitory synapses on the dendritic shaft (arrowheads in A) and in the spines (arrows in A).

### NKCC1a and NKCC1b are more mobile than KCC2 at the surface of dendrites

The clustering of NKCC1a suggests the involvement of the diffusion-capture mechanism in the regulation of the aggregates, as shown for KCC2 (Chamma et al., 2013; Heubl et al., 2017) and neurotransmitter receptors (Choquet and Triller, 2013; Triller and Choquet, 2018). Such mechanism would allow rapid control of the amount of transporters near the synapse and thus of chloride influx and homeostasis.

We first compared the membrane dynamics of NKCC1a and NKCC1b using Quantum Dot-based Single Particle Tracking (QD-SPT) technique in mature hippocampal neurons. To this end, neurons were transfected at DIV14 with recombinant HA-tagged NKCC1a or NKCC1b, stained live at 21–24 DIV with QD-coupled anti-HA antibody and real time imaging was performed (see Material and Methods). Then, NKCC1a and NKCC1b trajectories were reconstructed with homemade software and the diffusion coefficient and size of the explored area were extracted for each trajectory from the mean square displacement (MSD) plot versus time function. The observation of individual trajectories showed that NKCC1a and NKCC1b explore a large surface of the somato-dendritic membrane (Fig. 4A). Quantitative analysis performed on the whole (synaptic + extrasynaptic) population of trajectories revealed that the median value of the diffusion coefficient (Fig. 4B) and the median value of the explored area (Fig. 4C) were not significantly different between NKCC1a and NKCC1b, indicating similar diffusion properties. In comparison with KCC2, we found that NKCC1a and NKCC1b trajectories were longer and extended over a larger area of the membrane compared with those of KCC2 (Fig. 4A), suggesting **lower diffusion constraints for NKCC1a and NKCC1b in the plasma membrane**. This observation was corroborated by quantitative analysis showing increased diffusion coefficient (Fig. 4D) and greater explored area (Fig. 4E) for NKCC1a than KCC2 at the dendritic surface. Our data indicate that NKCC1a and NKCC1b are less restricted in their movement in the dendritic membrane than KCC2, likely reflecting **a weaker interaction of NKCC1a and NKCC1b with scaffolding molecules, which may result in weaker anchoring of NKCC1 transporters than KCC2.**

**Fig. 4.**
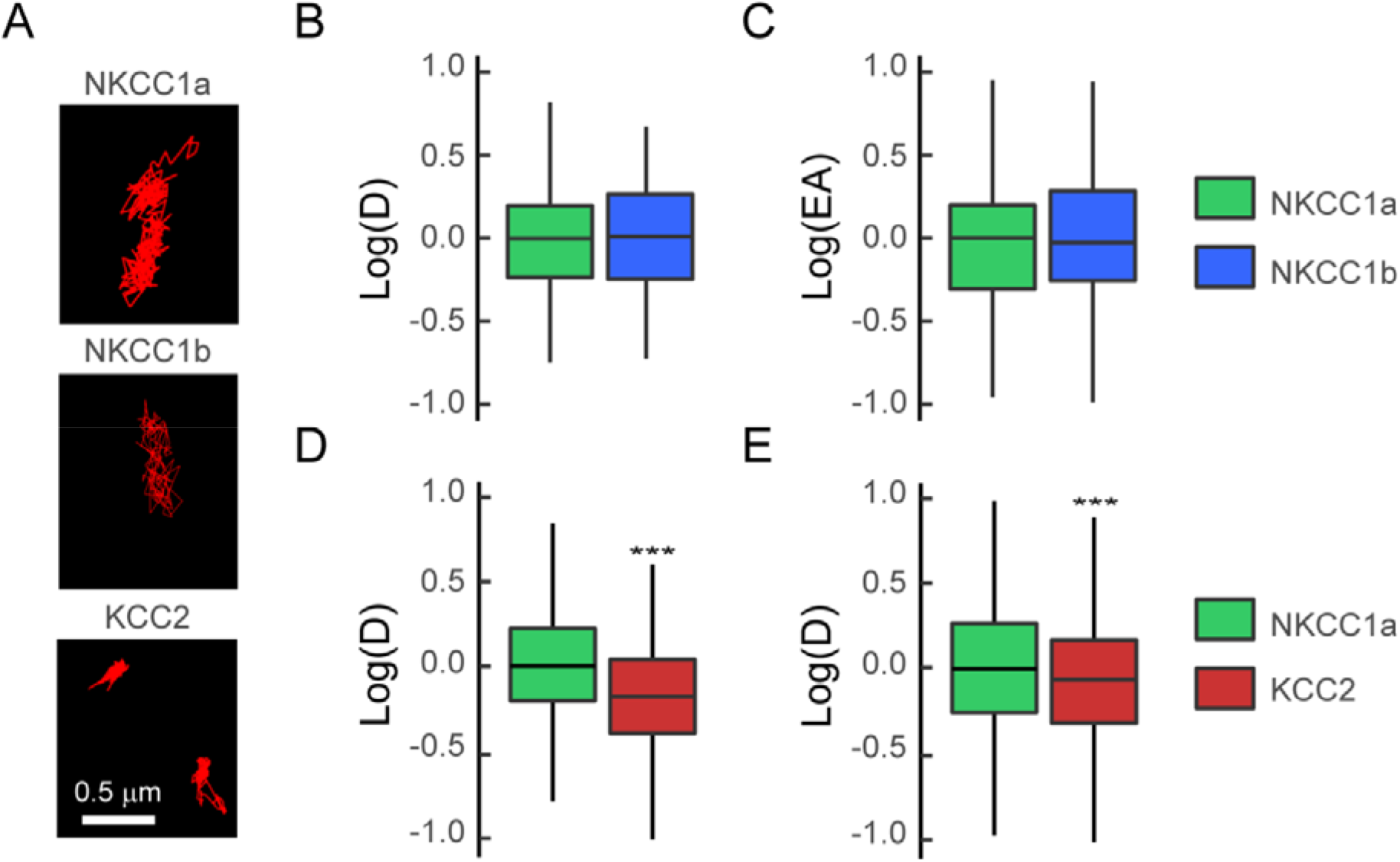
NKCC1a and NKCC1b are more mobile than KCC2 at the dendritic surface. **A**, Representative trajectories (red) of NKCC1a, NKCC1b and KCC2 at the surface of dendrites of mature 21-DIV old neurons. Scale bars: 0.5 μm. **B-C**, Similar log(D) (B) and log(EA) (C) for the bulk (extrasynaptic + synaptic) population of NKCC1a (green) and NKCC1b (blue). Diffusion coefficient (D): NKCC1a n = 146 QDs, NKCC1b n = 116 QDs, p = 0.19, 1 culture. Explored area (EA): p = 0.63. **D-E**, Reduced log(D) (D) and log(EA) (E) for the bulk population of KCC2 (red) as compared to NKCC1a (green). Diffusion coefficient (D): NKCC1a n = 180 QDs, KCC2 n = 675 QDs, p < 2.2 10^-16^, 3 cultures. Explored area (EA): p < 2.2 10^-16^. In B-E, data are presented as median values ± 25%–75% IQR. Values in B-C and D-E were normalized to the corresponding NKCC1a values. ***, p<1.0 10^-3^ (Welch t-test).

### NKCC1a and NKCC1b but not KCC2 are confined within endocytic zones

Since NKCC1a (Fig. 1 A-B) and NKCC1b (Fig. 1 C-D) are expressed at the dendritic membrane at a lower level than KCC2 (Fig. 1 E-F), we wondered if NKCC1a and NKCC1b have a particular relationship with endocytic zones. We analyzed the diffusion properties of NKCC1a and NKCC1b in endocytic zones identified by the presence of clathrin-YFP clusters in transfected neurons (Fig. 5A). Clathrin-YFP form numerous aggregates all along the dendrites with some of them being adjacent to GABA_A_R clusters (Fig. 5A), in agreement with the notion that GABAergic synapses are surrounded by clathrin-coated pits (Smith et al., 2012). Individual NKCC1a and NKCC1b trajectories exhibited decreased surface exploration in clathrin-YFP clusters compared to remotely located regions (Fig. 5B). This was consistent with the significant decrease in diffusion coefficients (Fig. 5C) and explored area (Fig. 5D) of NKCC1a in endocytic zones as compared to transporters detected at distance of clathrin-YFP clusters. Comparison of NKCC1a and NKCC1b revealed that they display similar values of diffusion coefficient (Fig. 5E) and surface exploration within (Fig. 5F) and outside (Fig. 5 G, H respectively) endocytic zones. Unlike NKCC1a and NKCC1b, the diffusion coefficient (Fig. 5I) and the size of the explored area (Fig. 5J) of KCC2 did not differ inside and outside clathrin-YFP clusters, indicating that, at least under basal activity condition, KCC2 is not confined within endocytic zones. We concluded that **NKCC1a and NKCC1b but not KCC2 chloride transporters are confined within endocytic zones under basal activity conditions**.

**Fig. 5.**
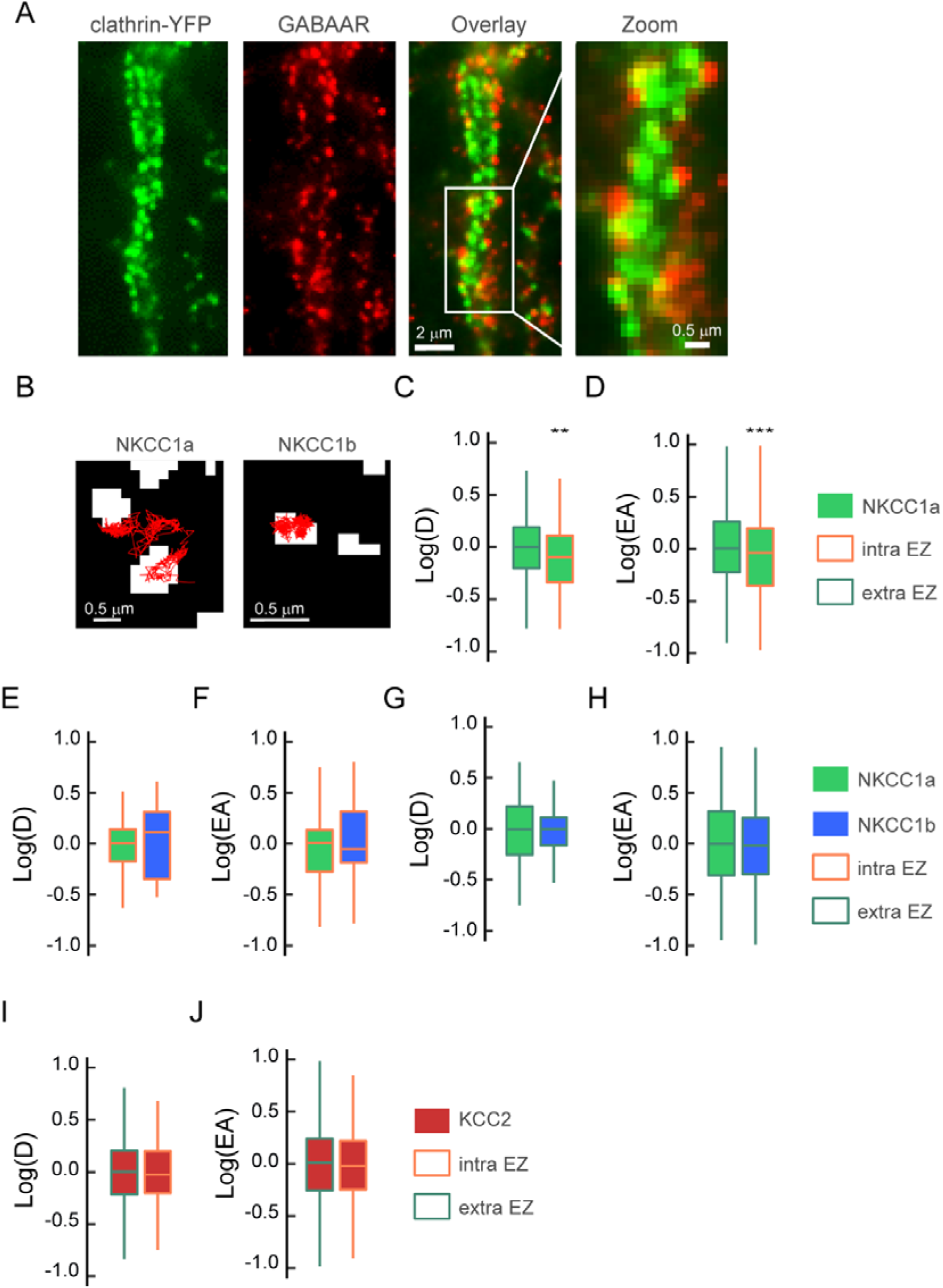
NKCC1a and NKCC1b but not KCC2 are confined within endocytic zones. **A**, Representative images of clathrin-YFP (green) and GABA_A_R γ2 subunit (red) in neurons transfected at DIV14 with clathrin-YFP and stained at DIV21 for the GABA_A_R γ2 subunit. Overlay shows that GABA_A_Rγ2 clusters are surrounded by clusters of clathrin-YFP. Scale bars: 2 μm and 0.5 μm as indicated. **B**, Representative trajectories (red) of NKCC1a and NKCC1b in relation with endocytic zones (white) identified by the presence of clathrin-YFP clusters. Scale bars: 0.5 μm. **C-D**, Significant reduction in log(D) (**C**) and log(EA) (**D**) for NKCC1a (green bars) trajectories inside (orange lines) endocytic zones as compared to QDs detected outside endocytic zones (green lines). Diffusion coefficient (D): intra EZ n = 353 QDs, extra EZ n = 180 QDs, p = 0.0013, 3 cultures. Explored area (EA): p = 1.65 10^-5^. **E-H**, Similar NKCC1a (green) and NKCC1b (blue) diffusion coefficient (**E, G**) and size of the explored area (**F, H**) inside (orange lines) or outside (green lines) endocytic zones. NKCC1a, intra EZ n = 64 QDs, extra EZ n = 84 QDs; NKCC1b, intra EZ n = 47 QDs, extra EZ n = 71 QDs, 1 culture. Diffusion coefficient (D): intra EZ p = 0.096; extra EZ p = 0.66. Explored area (EA): intra EZ p = 0.09; extra EZ p = 0.41. **I-J**, KCC2 transporters are not trapped in endocytic zones. No difference in log(D) (**J**) and log(EA) (**K**) for KCC2 (red) inside (orange lines) or outside (green lines) endocytic zones. Diffusion coefficient (D): intra EZ n = 306 QDs, extra EZ n = 675 QDs, p = 0.35, 3 cultures. Explored area (EA): KCC2, p = 0.077. In all graphs, data are presented as median values ± 25%–75% IQR. In C-D, NKCC1a values in EZ were normalized to extra EZ values. In E-F and G-H, NKCC1b values intra EZ or extra EZ were normalized to NKCC1a intra EZ or extra EZ values, respectively. In I-J, KCC2 intra EZ values were normalized to extra EZ values. **, p<1.0 10^-2^, ***, p<1.0 10^-3^ (Welch t-test).

### The lateral diffusion of NKCC1a is tuned by glutamatergic activity in the axon but not in the dendrite

Since we have previously demonstrated that a rapid regulation of KCC2 diffusion by excitatory glutamatergic and inhibitory GABAergic activities tune the membrane stability of KCC2 and thereby chloride homeostasis (Chamma et al., 2013; Heubl et al., 2017), we asked whether NKCC1a membrane dynamics in the axon and in the dendrites could also be modulated by glutamatergic activity. To this end, NKCC1a diffusion was explored in the presence of the Na^+^ channel blocker tetrodotoxin TTX (1μM), the ionotropic glutamate receptor antagonist kynurenic acid (KYN, 1 mM), and the group I/group II mGluR antagonist R,SMCPG (500 μM) (as done in Heubl et al., 2017). Cells were exposed to the cocktail “TTX + KYN + MCPG” 10 minutes before recording and were imaged up to one hour in the presence of the drugs. We observed that, when exposed to TTX + KYN + MCPG, exploration of individual QDs was limited to smaller areas in the axon (Fig 6 A-C), compared to QDs on dendrites (Fig. 6 D-F). Statistical analysis of hundreds of trajectories showed that the slope of the mean squared displacement (MSD) as a function of time was reduced for trajectories recorded in the axon compared to those recorded in the dendrite (Fig. 6G), indicating increased confinement of NKCC1a in the axon. Consistent with this observation, the median value of the diffusion coefficient (Fig. 6H) was significantly lower in the axon compared to the dendrite. The cumulative log(D) graph describes an overall trend with no apparent sub-populations of QDs (Fig. 6I). The size of the explored area (Fig. 6J) was smaller in the axon compared to the dendrite and the cumulative probability curve showed that all trajectories behaved similarly (Fig. 6K). Therefore, NKCC1a confinement is higher in the axon than in the dendrite in conditions of glutamatergic activity blockade. These results contrast with those obtained in the absence of activity blockers showing reduced confinement in the axon vs. the dendrite (Côme et al., 2019b). Indeed, when directly comparing NKCC1a lateral diffusion in the absence or presence of TTX + KYN + MCPG, the diffusion coefficient and explored area were significantly reduced upon drug treatment in the axon while it was unchanged in the dendrite (Fig. 6 L, M, respectively). Therefore, we concluded that **NKCC1a diffusion is reduced in the axon but not in the dendrite upon glutamatergic activity blockade**.

**Fig. 6.**
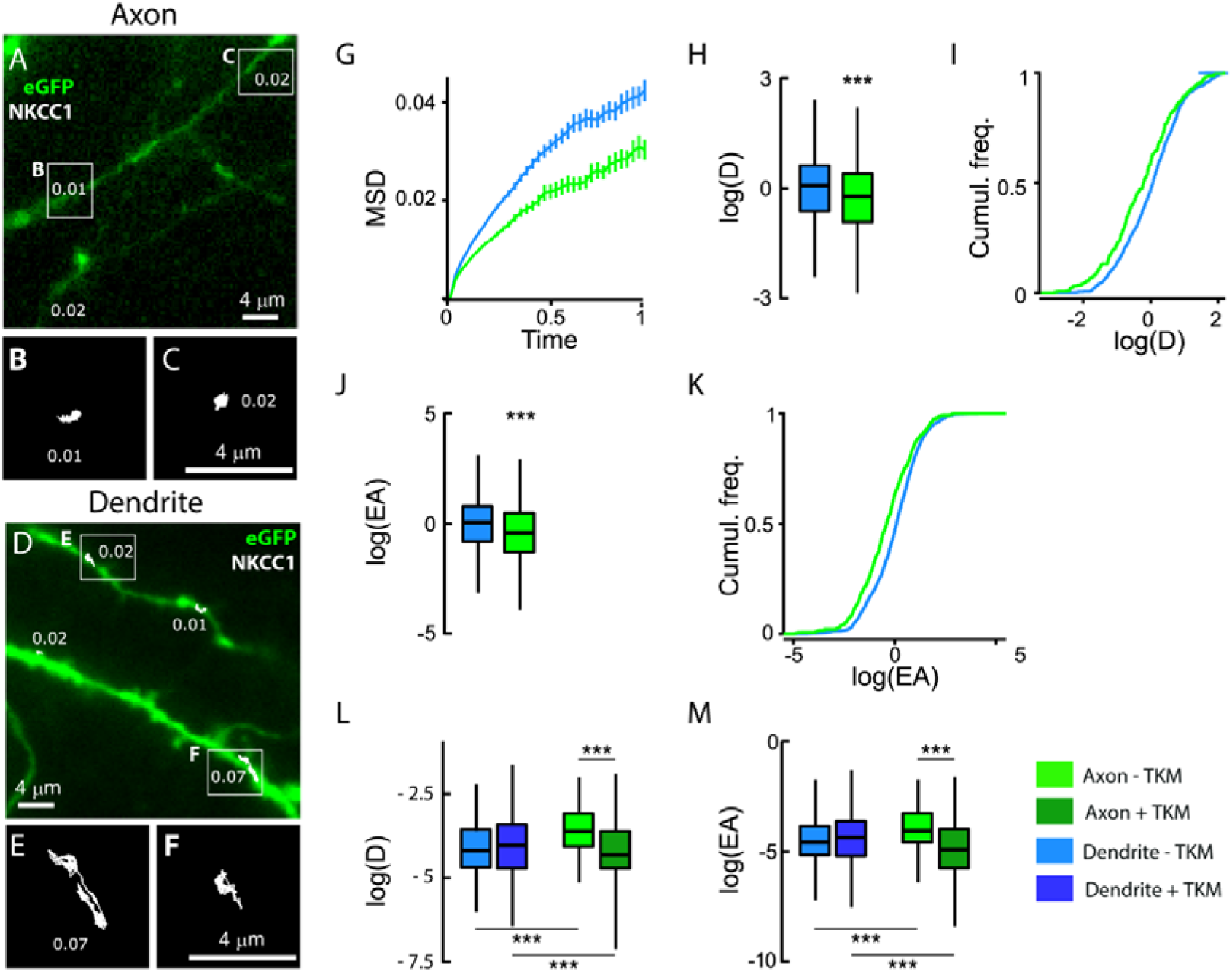
Increased confinement of NKCC1a in the axon but not in the dendrite upon glutamatergic activity blockade. **A-F**, Representative trajectories (white in A-F) of QD-bound HA-tagged recombinant NKCC1a along the axon (A-C), and on dendrites (D-F) of neurons treated 10 min to 1hour with TTX + KYN + MCPG (TKM). QD trajectories (white in A-F) were overlaid with fluorescent labeling of cytoplasmic eGFP to label neurites (green in A, D). Mean diffusion coefficient values (μm^2^.s^-1^) are displayed for each trajectory. Scale bars: 4 μm. Note that NKCC1a trajectories are shorter in the axon than in the dendrite suggesting increased confinement in the axon. **G**, Time-averaged MSD functions of dendritic QDs (blue) and axonal QDs (green). The MSD versus time relationship for dendritic trajectories shows a steeper initial slope, suggesting that trajectories are less confined. Dendrite, n = 707 QDs, Axon, n = 305 QDs, 4 cultures. **H-I**, Box plots (H) and cumulative probabilities (I) of QD diffusion coefficients in dendritic (blue) or axonal (green) compartments. Note the reduced diffusion in the axon. p = 6.3 10^-4^. **J-K**, Box plots (J) and cumulative probabilities (K) of explored area EA in dendritic (blue) or axonal (green) compartments. Note the reduced exploration in the axon. p = 2.3 10^-8^. **L-M**, Comparison of NKCC1a diffusion coefficient (L) and surface explored (M) in absence (-TKM) or presence of TTX + KYN + MCPG (+TKM), showing reduced speed and increased confinement for axonal but not dendritic QDs upon TKM exposure. No TKM: N_dendrite_ = 525 QDs, N_axon_ = 117 QDs. +TKM: N_dendrite_ = 2010 QDs, N_axon_ = 305 QDs, 3 cultures. Diffusion coefficient (D): Dendrite, p < 1.0 10^-3^ Axon, p = 8.4 10^-13^. Explored area (EA): Dendrite, p < 1.0 10^-3^; Axon, p = 8.4 10^-13^. In H, J, L, M, data are presented as median values ± 25%–75% IQR. In H, J, NKCC1a values in axons were normalized to values obtained in dendrites. ***, p<1.0 10^-3^ (Welch t-test).

We then analyzed the effect of TTX + KYN + MCPG treatment on the diffusion properties of NKCC1a in relation to synapses. For this, we studied the diffusive behavior of NKCC1a on the dendrites of neurons co-transfected with NKCC1a-HA and the inhibitory and excitatory synaptic markers Gephyrin-Finger-YFP (Fig. 7 A-C, G) and Homer1c-DsRed, respectively (Fig. 7 D-F, G). We found that individual QDs explored a restricted area of the membrane near GABAergic synapses (Fig. 7 A-C, G) and glutamatergic synapses (Fig. 7 D-F, G) compared with extrasynaptic regions (Fig. 7G). However, we observed that NKCC1a transporters sometimes escaped from the perisynaptic region to explore neighboring synapses whether they were excitatory or inhibitory synapses (Fig. 7G). This is reminiscent of the diffusive behavior of KCC2 (Chamma et al., 2013; Heubl et al., 2017).

**Fig. 7.**
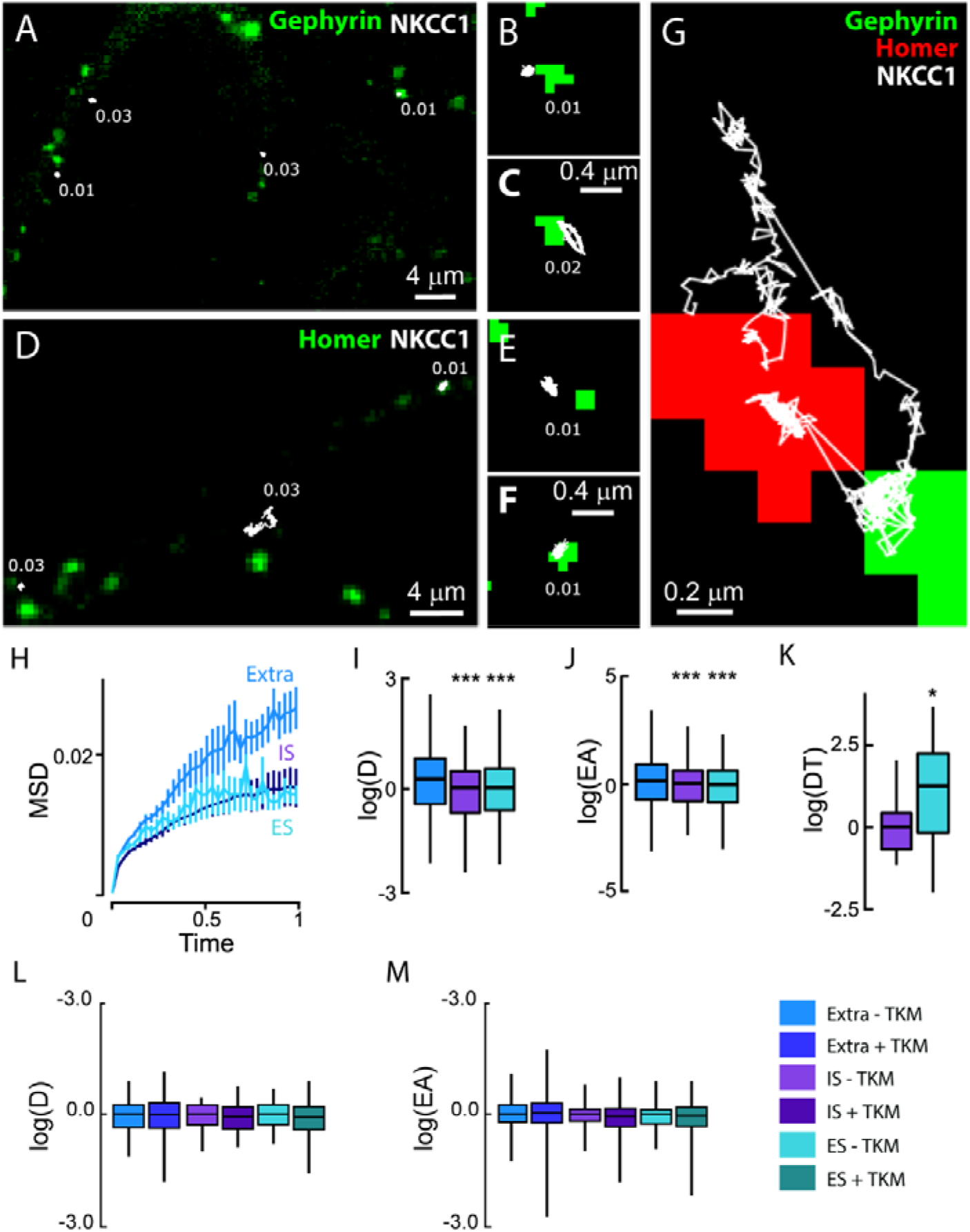
No effect of glutamatergic activity blockade on NKCC1a diffusion at synaptic and extrasynaptic sites. **A-F**, Representative trajectories (white) of QD-bound HA-tagged NKCC1a at the surface of gephyrin-Finger-YFP (A-C) or homer1c-DsRed (D-F) transfected neurons. QD trajectories (white) were overlaid with fluorescent clusters of Gephyrin-Finger-YFP (green in A-C), or Homer1c-DsRed (green in D-F) to identify inhibitory (IS) and excitatory (ES) synapses, respectively. Scale bars: 4 μm in A and D and 0.4 μm in B-C and E-F. **G**, An individual NKCC1a transporter shifts between an excitatory (green) and an inhibitory synapse (red). Scale bar: 0.2 μm. **H**, Time-averaged MSD functions of extrasynaptic QDs (blue), QDs at excitatory synapses (turquoise), and QDs at inhibitory synapses (purple). The MSD versus time relationship for extrasynaptic trajectories shows a steeper initial slope, suggesting that trajectories were less confined. Extra, n = 747 QDs, IS, n = 209 QDs, ES, n = 369 QDs, 5 cultures. **I-J**, Boxplots of log(D) (I) and log(EA) (J) showing a reduced mobility and an increased confinement of NKCC1a at IS (purple) and ES (turquoise) compared to transporters located at distance of synapses (blue). Diffusion coefficient (D): IS, p = 1.58 10^-14^; ES, p = 8.4 10^-13^. Explored area (EA): IS, p = 5.6 10^-11^; ES, p = 7.2 10^-10^. **K**, Boxplots of log(dwell time, DT) of NKCC1a at inhibitory (purple) and excitatory (turquoise) synapses showing increased DT at excitatory synapses as compared with inhibitory synapses. p = 0.029. **L-M**, Boxplots of log(D) (L) and log(EA) (M) of NKCC1a at extrasynaptic sites (blue), IS (purple) and ES (turquoise) in absence (light color) or presence (dark color) of TKM. No TKM: Extra, n = 152 QDs, IS, n = 17 QDs, ES, n = 46 QDs; TKM: Extra, n = 1545 QDs, IS, n =311 QDs, ES, n = 368 QDs, 2-3 cultures. Diffusion coefficient (D): Extra, p = 0.31; IS, p = 0.39; ES, p = 0.998. Explored area (EA): Extra, p = 0.428; IS, p = 0.021; ES, p = 0.015. In I-M, data are presented as median values ± 25%–75% IQR. In I, J, NKCC1a synaptic values were normalized to extrasynaptic values (all in -TKM). In K, values at ES were normalized to values obtained at IS (all in –TKM). In L, M, Values obtained +TKM were compared to values -TKM. *, p<5.0 10^-2^, ***, p<1.0 10^-3^. (I, J, L, M, Welch t-test, K, Mann–Whitney rank sum test).

We next measured the synaptic and extrasynaptic mobility of NKCC1a on hundreds of trajectories. For receptors enriched in the postsynaptic membrane (e.g. GABA_A_R, glycine receptor or AMPAR…), trajectories are usually considered synaptic when they overlap the synaptic mask or extrasynaptic for spots located two pixels (380 nm) apart (Chamma et al., 2013; Heubl et al., 2017). Here, the application of this parameter did not detect enough synaptic trajectories of NKCC1a. Having shown using super-resolution imaging that NKCC1a form perisynaptic nanoclusters (Fig. 3 A-C), we assumed that some NKCC1a transporters should be confined near synapses. We therefore progressively increased the area around the synaptic mask until we identified a zone that separated confined and free trajectories. Thus, a threshold of 4 pixels (760 nm) was determined. This is in agreement with a perisynaptic localization of the NKCC1 transporters. However, this size is twice larger than the one required to separate the confined vs. freely diffusing KCC2 transporters (Chamma et al., 2013; Heubl et al., 2017). **This suggests that KCC2 is distributed as a perisynaptic ring itself surrounded by a ring of NKCC1.**

We report a reduced slope of the MSD vs. time function for perisynaptic trajectories as compared with extrasynaptic ones (Fig. 7H), indicative of increased confinement of NKCC1a at the periphery of both excitatory and inhibitory synapses. Consistently, the median values of the diffusion coefficient and explored area were also significantly decreased at the periphery of glutamatergic and GABAergic synapses (Fig. 7 I, J respectively). Therefore, **NKCC1a is confined at the periphery of synapses, regardless of whether (this work) or not (Côme et al., 2019b) glutamatergic transmission is blocked.** Although the diffusion coefficient and the size of the explored area of NKCC1a were similar near both excitatory and inhibitory synapses, the synaptic dwell time of NKCC1a near glutamatergic synapses was ~1.9 fold that of the transporter near inhibitory synapses (Fig. 7K). Thus, we concluded that **NKCC1a is more confined near glutamatergic synapses than near GABAergic synapses**. These results obtained in conditions of glutamatergic activity blockade do not differ from those obtained in conditions that preserved neuronal activity (Côme et al., 2019b). Analysis of TTX + KYN + MCPG treatment revealed no effect of blocking glutamatergic activity on the diffusion coefficient (Fig. 7L) and the size of the explored zone (Fig. 7M) at inhibitory synapses, excitatory synapses or in the extrasynaptic membrane. **The lack of regulation by glutamatergic activity of NKCC1a lateral diffusion in the dendrite (inside and outside synapses) supports our hypothesis that NKCC1a is regulated in the axon but not in the dendrite by glutamatergic activity.**

## Discussion

We have studied, in mature hippocampal neurons, the diffusion and the clustering of the co-transporters NKCC1a and NKCC1b. First, we showed that both transporters display a heterogeneous distribution at the plasma membrane: NKCC1a and NKCC1b are either diffusely distributed in the axonal and somato-dendritic membranes or are organized in nanoclusters. In the dendrite, NKCC1a accumulates in the near vicinity of inhibitory synapses formed on the dendritic shaft where the transporter is slowed down and confined. NKCC1a and NKCC1b transporters are also confined in endocytic zones in the dendrite. To be organized in membrane clusters, NKCC1a and NKCC1b transporters are probably anchored to the cytoskeleton via their binding to yet undefined scaffolding proteins. Future studies will tell whether distinct scaffolding molecules anchor NKCC1a and NKCC1b in the dendritic vs. the axon. However, the binding of NKCC1a and NKCC1b to the cytoskeleton is probably weaker than that anchoring KCC2 to the cytoskeleton since we show here using STORM that NKCC1a and NKCC1b clusters are half the size and are significantly less dense in molecules than KCC2 clusters. These different clustering properties were associated with the fact that NKCC1a and NKCC1b are much more mobile than KCC2 on the neuronal surface.

Acute blockade of basal glutamatergic activity with TTX+KYN+MCPG showed that only the lateral diffusion of NKCC1 in the axon was rapidly modified by the treatment while the mobility and confinement of the dendritic transporter was not changed whatever the synaptic or extrasynaptic compartment analyzed. This reports that the regulation of the membrane dynamics of NKCC1a is different in the axon *vs*. dendrite and may involve separate regulatory mechanisms. Further studies will be needed to identify the mechanism of axonal regulation of NKCC1a diffusion and the role of increased NKCC1a confinement in the axon during reduced basal glutamatergic activity. This could be potentially important in the regulation of action potential firing in mature pyramidal cells and thus of synaptic glutamate release (Khirug et al., 2008). Chandelier interneurons establish axo-axonal GABAergic synapses at the AIS of pyramidal cells (Khirug et al., 2008). The presence of NKCC1 in the AIS maintains a depolarized E_GABA_ value (Khirug et al., 2008). By exciting presynaptic GABA_A_R, GABA release from these interneurons would facilitate spontaneous glutamate release from presynaptic terminals (Jang et al., 2006). Indeed, interneurons depolarize or even excite mature cortical pyramidal neurons (Szabadics et al., 2006; Woodruff et al., 2009). The regulation of NKCC1 diffusion-capture in the axon by glutamatergic activity would be a way to control glutamate release at synaptic terminals.

One hypothesis is that increased glutamatergic activity would increase diffusion of NKCC1 along the axon to decrease its clustering and activity along the axon, thereby limiting its contribution to membrane depolarization. This would therefore correspond to a homeostatic mechanism controlling the discharge of action potentials and the release of glutamate at presynaptic sites. This supports the hypothesis of an essential modulatory function of NKCC1 on glutamate release at photoreceptor synaptic terminals in the tiger salamander retina (Shen et al., 2013). The fact that this regulation predominates in the axon and not in the dendrite highlights the primary role of NKCC1 in the axon at least under physiological conditions when its membrane distribution is maintained at a low level in the dendrite.

Although this pathway has little influence on the amount/function of NKCC1 at the neuronal dendritic surface under basal activity conditions, it may play a role in pathological situations associated with increased expression of NKCC1 (Kourdougli et al., 2017). Endocytic zones have been shown to constitute reserve pools of molecules, which can, depending on the synaptic demand, be released and reintegrated into the diffusing pool of receptors (Petrini et al., 2009). In the case of NKCC1, this reserve pool would allow a rapid increase in the transporter availability in the dendritic plasma membrane, e.g. in pathological situations in which an up-regulation of NKCC1 has been observed (Kourdougli et al., 2017). In the pathology, upregulation of NKCC1 is often accompanied by a down-regulation of KCC2 at the neuronal surface (Liu et al., 2020; Puskarjov et al., 2012). KCC2 is also regulated by diffusion-capture. We have shown that an acute exposure of neurons to the convulsive agent 4-AP increases the lateral diffusion of KCC2 transporters, which escape the clusters and are internalized and degraded (Chamma et al., 2013). Thus, lateral diffusion would be a general mechanism to control the membrane stability of the chloride co-transporters NKCC1 and KCC2. Inhibition of the pathway controlling the lateral diffusion of NKCC1 and/or KCC2 would prevent the loss of KCC2 and the increase in NKCC1 at the surface of the neuron, thus preventing the abnormal rise of [Cl^-^]_i_ in the pathology and the resulting adverse effects.

## Limitations of study

All results were used for analysis except in few cases. Cells with signs of suffering (apparition of blobs, fragmented neurites) were discarded from the analysis.

## Acknowledgments

We thank J. Nabekura for kindly providing the original pEGFP–IRES–KCC2 full-length construct, D. Choquet for the homer1c–GFP construct. We are grateful to the Cell and Tissue Imaging Facility of Institut du Fer à Moulin (IFM). This work was supported by Institut National de la Santé et de la Recherche Médicale, Sorbonne Université-UPMC as well as by the Agence Nationale de la Recherche (ANR WATT ANR-19-CE16-0005), Fondation pour la Recherche sur le Cerveau, Fondation Française pour la Recherche sur l’Épilepsie and Fondation pour la Recherche Médicale. EC and EP are the recipients of a doctoral fellowship from the Sorbonne Université and EC was a recipient of a 4-year PhD grant from the Fondation pour la Recherche Médicale. The STORM/PALM microscope was supported by DIM NeRF from Région Ile-de-France and by the FRC/Rotary ‘Espoir en tête’.

## Author contributions

SL conceptualized the project, designed the experiments and supervised the experimental work. SL and EC prepared the figures and SL wrote the paper. EC and JG performed single particle tracking and analyzed the data. EC, JG and EP performed conventional wide field microscopy and analyzed the data. EC, ZM and XM performed super-resolution imaging and EC and JG analyzed the data. SSL performed confocal imaging. MR prepared the hippocampal cultures. IM raised the NKCC1 constructs.

## Data and materials availability

The data that support the findings of this study are available from the corresponding author upon reasonable request.

## RESOURCE AVAILABILITY

### Lead contact

Any additional information or enquiries about reagents and resources should be directed to the Lead contact, Sabine Lévi

### Materials availability

The transfer of plasmids generated for this study will be made available upon request. A Materials Transfer Agreement may be required.

### Data and code availability

No standardized datatypes are reported in this paper. All data reported in this paper will be shared by the lead contact upon request. This paper does not report original code. Any additional information required to reanalyze the data reported in this paper is available from the lead contact upon request.

## EXPERIMENTAL MODEL AND SUBJECT DETAILS

For all experiments performed on primary cultures of hippocampal neurons, animal procedures were carried out according to the European Community Council directive of 24 November 1986 (86/609/EEC), the guidelines of the French Ministry of Agriculture and the Direction Départementale de la Protection des Populations de Paris (Institut du Fer à Moulin, Animalerie des Rongeurs, license C 72-05-22). All efforts were made to minimize animal suffering and to reduce the number of animals used. Timed pregnant Sprague-Dawley rats were supplied by Janvier Lab and embryos were used at embryonic day 18 or 19 as described below.

## Material and Methods

### Dissociated hippocampal cultures

Primary cultures of hippocampal neurons were prepared as previously described (Chamma et al. 2013) with some modifications in the protocol. Briefly, hippocampi were dissected from embryonic day 18 or 19 Sprague-Dawley rats of either sex. Tissue was then trypsinized (0.25% v/v), and mechanically dissociated in 1× HBSS (Invitrogen, Cergy Pontoise, France) containing 10mM HEPES (Invitrogen). Neurons were plated at a density of 120 × 103 cells/ml onto 18-mm diameter glass coverslips (Assistent, Winigor, Germany) pre-coated with 50 μg/ml poly-D,Lornithine (Sigma-Aldrich, Lyon, France) in plating medium composed of Minimum Essential Medium (MEM, Sigma) supplemented with horse serum (10% v/v, Invitrogen), L-glutamine (2 mM) and Na+ pyruvate (1 mM) (Invitrogen). After attachment for 3–4 h, cells were incubated in culture medium that consists of Neurobasal medium supplemented with B27 (1X), L-glutamine (2 mM), and antibiotics (penicillin 200 units/ml, streptomycin, 200 μg/ml) (Invitrogen) for up to 4 weeks at 37 °C in a 5% CO_2_ humidified incubator. Each week, one fifth of the culture medium volume was renewed.

## METHOD DETAILS

### DNA constructs

The pcDNA3.1 Flag YFP hNKCC1 HA-ECL2 (NT931) was a gift from Biff Forbush (Addgene plasmid # 49063; http://n2t.net/addgene:49063; RRID:Addgene_49063; (Somasekharan and Forbush 2013). From this NKCC1-HA-Flag-mVenus plasmid, the following constructs were raised: NKCC1-HA-Δflag-ΔmVenus by truncation of the tags located on NKCC1 NTD. The following constructs were also used: pCAG_rat KCC2-3Flag-ECL2 (Chamma et al. 2013), eGFP (Clontech), pCAG_GPHN.FingR-eGFP-CCR5TC (Gross et al. 2013) (gift from Don Arnold, Addgene plasmid # 46296; http://n2t.net/addgene:46296; RRID:Addgene_46296), EYFP-Clathrin (Rust et al., 2004) was a gift from Xiaowei Zhuang (Addgene plasmid # 20921; http://n2t.net/addgene:20921; RRID:Addgene_20921), homer1c-DsRed (kindly provided by D. Choquet, IIN, Bordeaux, France). All constructs were sequenced by Beckman Coulter Genomics (Hope End, Takeley, U.K).

### Neuronal transfection

Neuronal transfections were carried out at DIV 13–14 using Transfectin (BioRad, Hercules, USA), according to the manufacturers’ instructions (DNA:transfectin ratio 1 μg:3 μl), with 1–2 μg of plasmid DNA per 20 mm well. Simple transfections of NKCC1-HA-Flag-mVenus or KCC2-Flag were done with a plasmid concentration of 1 μg. The following ratios of plasmid DNA were used in co-transfection experiments: 1:0.4:0.4 μg for NKCC1 constructs together with GPHN.FingR-eGFP and homer1c-DsRed; 1:0.2 μg for NKCC1 constructs with eGFP; 0.7:0.7 μg for NKCC1 or KCC2 constructs with gephyrin-dendra2. Experiments were performed 7–10 days posttransfection. SPT, STORM, and STORM/PALM experiments were performed with Δflag-ΔmVenus NKCC1 constructs. Standard epifluorescence microscopy was done with Flag-mVenus NKCC1 constructs.

### Pharmacology

The following drugs were used: TTX (1 μM; Latoxan, Valence, France), R,S-MCPG (500 μM; Abcam, Cambridge, UK), S-MCPG (250 μM; HelloBio), Kynurenic acid (1 mM; Abcam). R,S-MCPG and S-MCPG were prepared in equimolar concentrations of NaOH; TTX in 2% citric acid (v/v); closantel in DMSO (Sigma). Equimolar DMSO concentrations were used for control experiments in these conditions. For SPT experiments, neurons were transferred to a recording chamber, pre-incubated in presence of drugs and/or peptide at 31 °C for 10 min in imaging medium (see below for composition) and used within 45 min in presence of the appropriate drug for imaging. For immunofluorescence experiments, drugs were added directly to the culture medium for 30 min in a CO_2_ incubator set at 37 °C. The imaging medium consisted of phenol red-free minimal essential medium supplemented with glucose (33 mM; Sigma) and HEPES (20 mM), glutamine (2 mM), Na^+^-pyruvate (1 mM), and B27 (1X) from Invitrogen.

### Live cell staining for single-particle imaging

Neurons were stained as described previously (Bannai et al., 2006). Briefly, cells were incubated for 3-8 min at 37 °C with primary antibodies against HA (rabbit, 1:250, Cell signaling Technology, cat #C29F4) or Flag (mouse, 1:700, Sigma, cat #F3165) for NKCC1 and KCC2 labeling, respectively. After washes, cells for NKCC1 detetcion were incubated for 1 min with F(ab’)2-Goat anti-Rabbit IgG (H+L) Secondary Antibody QDot emitting at 655 nm (1 nM; Invitrogen) in PBS (1 M; Invitrogen) supplemented with 10% Casein (v/v) (Sigma). Cells for KCC2 detection were incubated for 3-5 min at 37°C with biotinylated Fab secondary antibodies (goat antimouse: 1:700; Jackson Immuno research, cat #115-067-003, West Grove, USA) in imaging medium. After washes, cells were incubated for 1 min with streptavidin-coated quantum dots (QDs) emitting at 605 nm (1 nM; Invitrogen) in borate buffer (50 mM) supplemented with sucrose (200 mM) or in PBS (1 M; Invitrogen) supplemented with 10% Casein (v/v) (Sigma).

### Single-particle tracking and analysis

Cells were imaged as previously described using an Olympus IX71 inverted microscope equipped with a 60X objective (NA 1.42; Olympus) and a 120W Mercury lamp (X-Cite 120Q, Lumen Dynamics). Individual images of gephyrin-YFP and homer1c-DsRed, and QD real time recordings (integration time of 30 ms over 1200 consecutive frames) were acquired with an ImagEM EMCCD camera and MetaView software (Meta Imaging 7.7). Cells were imaged within 45 min following appropriate drugs pre-incubation. QD tracking and trajectory reconstruction were performed with homemade software (Matlab; The Mathworks, Natick, MA) as described in (Bannai et al., 2006). One to two sub-regions of dendrites were quantified per cell. In cases of QD crossing, the trajectories were discarded from analysis. Trajectories were considered synaptic when overlapping with the synaptic mask of gephyrin-mRFP or homer1c-GFP clusters, or extrasynaptic for spots two pixels (380 nm) or four pixels (760 nm) away for KCC2 and NKCC1, respectively. The inclusion area was increased for NKCC1 relatively to previous studies on KCC2 (Chamma et al., 2013; Heubl et al., 2017) due to the low numbers of NKCC1 trajectories recorded with a 380 nm distance. Expanding the radius did actually not change results for perisynaptic NKCC1 lateral diffusion, suggesting its clusters are located further away from synapses than KCC2 ones. Values of the mean square displacement (MSD) plot vs. time were calculated for each trajectory by applying the relation:

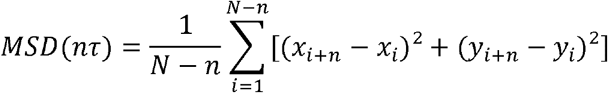

where τ is the acquisition time, N is the total number of frames, n and i are positive integers with n determining the time increment. Diffusion coefficients (D) were calculated by fitting the first four points without origin of the MSD vs. time curves with the equation: MSD(nτ) = 4Dnτ + σ; where σ is the spot localization accuracy. Depending on the type of lamp used for imaging, the QD pointing accuracy is ~20–30 nm, a value well below the measured explored areas (at least 1 log difference). The explored area of each trajectory was defined as the MSD value of the trajectory at two different time intervals of 0.42 and 0.45 s (Renner et al., 2012). Synaptic dwell time was defined as the duration of detection of QDs at synapses on a recording divided by the number of exits as detailed previously (Bannai et al., 2006). Dwell times ≤ 5 frames were not retained. The number of QD vary from one cell to another and from one condition to another in a given experiment.

### Immunocytochemistry

NKCC1-HA-Flag-mVenus membrane expression and clustering was assessed with staining performed after a 4-5 min fixation at room temperature (RT) in paraformaldehyde (PFA; 4% w/v; Sigma) and sucrose (20% w/v; Sigma) solution in 1× PBS. The cells were then washed in PBS and incubated for 30 min at RT in goat serum (GS; 3% v/v; Invitrogen) in PBS to block non-specific staining. Neurons were then incubated for 60-180 min at RT with HA antibody (rabbit, 1:250, Cell signaling Technology, cat #C29F4) in PBS–GS blocking solution. After washing, neurons were incubated with Cy™3 AffiniPure Donkey Anti-Rabbit IgG (H+L) (1.9 μg/ml; Jackson ImmunoResearch, cat #111-165-003) for standard epifluorescence assays, or Alexa Fluor^®^ 647 AffiniPure Donkey Anti-Rabbit IgG (H+L) (2 μg/ml, Jackson ImmunoResearch, cat #711-605-152) for super-resolution experiments, in PBS-GS solution. The coverslips were then washed, and mounted on slides with mowiol 844 (48 mg/ml; Sigma).

KCC2-Flag membrane clustering was assessed with live cell staining. Pre-treated neurons expressing KCC2-Flag were washed in imaging medium and incubated for 20 min at 4 °C with mouse primary antibody against Flag (1:400; Sigma, cat #F3165) in imaging medium. After washes with imaging medium, cells were fixed for 15 min at RT in 4% PFA and 20% sucrose solution in 1× PBS. The cells were then washed in PBS and incubated for 30 min at RT in goat serum (GS; 20% v/v; Invitrogen) in PBS to block non-specific staining. Neurons were then incubated for 45 min at RT with Cy5-conjugated goat anti-mouse antibodies (1.9 g/ml; Jackson Immuno Research, cat#115-175-205) in PBS–GS blocking solution, washed, and mounted on slides.

## QUANTIFICATION AND STATISTICAL ANALYSIS

### Fluorescence image acquisition and analysis

Sets of neurons compared for quantification were labeled and imaged simultaneously. Image acquisition was performed using a ×100 objective (NA 1.40) on a Leica (Nussloch, Germany) DM6000 upright epifluorescence microscope with a 12-bit cooled CCD camera (Micromax, Roper Scientific) run by MetaMorph software (Roper Scientific, Evry, France). Quantification was performed using MetaMorph software (Roper Scientific). To assess NKCC1 and KCC2 clusters, exposure time was fixed at a non-saturating level and kept unchanged between cells and conditions. For the dendritic intensity and clustering analysis, the ROI was precisely traced around focused dendrites, and global ROI pixel intensity was measured. For the clustering images were flatten background filtered (kernel size, 3 × 3 × 2) to enhance cluster outlines, and a user defined intensity threshold was applied to select clusters and avoid their coalescence. Clusters were outlined and the corresponding regions were transferred onto raw images to determine the mean NKCC1 or KCC2 cluster number, area and fluorescence intensity. The dendritic surface area of the region of interest was measured to determine the number of clusters per pixel. For each culture, we analysed ~10-15 cells per experimental condition.

### STORM and PALM microscopy

Cells were transfected at DIV14 with dendra2-gephyrin construct. They were then fixed at DIV21 in 4% PFA for 15 min, washed in PBS 1X and stained for KCC2 or NKCC1. They were then mounted in a Ludin chamber and imaged in PBS 1X. STORM and PALM imaging on fixed samples was conducted on an inverted N-STORM Nikon Eclipse Ti microscope with a 100× oil immersion objective (NA 1.49) and an Andor iXon Ultra EMCCD camera (image pixel size, 160 nm), using specific lasers for STORM imaging of Alexa 647 (640 nm) and for PALM imaging of dendra2 and mEos2 (405 and 561 nm). KCC2 or NKCC1 videos of of 30,000 frames were acquired at frame rates of 50 ms. Gephyrin-dendra2 movies of ~20000 frames were acquired at frame rates of 20 ms. The z position was maintained during acquisition by a Nikon perfect focus system. Single-molecule localization and 2D image reconstruction was conducted as described in Specht et al. (2013) by fitting the PSF of spatially separated fluorophores to a 2D Gaussian distribution. The position of fluorophore were corrected by the relative movement of the synaptic cluster by calculating the center of mass of the cluster throughout the acquisition using a partial reconstruction of 2000 frames with a sliding window (Specht et al., 2013). STORM and PALM images were rendered by superimposing the coordinates of single molecule detections, which were represented with 2D Gaussian curves of unitary intensity. To correct multiple detections coming from the same molecule, we identified detections occurring in the vicinity of space (2σ) and time (15 s) as belonging to the same molecule. The surface of NKCC1 and KCC2 clusters and the densities of molecules per square nanometer were measured in reconstructed 2D images through cluster segmentation based on detection densities. The minimal thresholds to determine clusters were 1% intensity, 0.1 per nm^2^ minimum detection density and 10 detections. The resulting binary image was analyzed with the function “regionprops” of Matlab to extract the surface area of each cluster identified by this function. Density was calculated as the total number of detections in the pixels (pixel size = 20 nm) belonging to a given cluster, divided by the area of the cluster.

### Statistics

Sampling corresponds to the number of quantum dots for SPT and number of cells for ICC. Sample size selection for experiments was based on published experiments, pilot studies, as well as in-house expertise. All results were used for analysis except in few cases. For imaging experiments (SPT, immunofluorescence), cells with signs of suffering (apparition of blobs, fragmented neurites) were discarded from the analysis. Data representation was usually done with boxplots or cumulative frequency plots. The statistical test to compare two groups was either Welch t-test when normality assumption was met (Q-Q plots and cumulative frequency fit), otherwise Mann-Whitney test was performed to assess the presence of a dominance or no between the two distributions. For variables following a log-normal distribution, such as variables obtained from SPT and STORM assays, we applied the log(.) function after division by the control group’s median value. For super-resolution experiments, as an important variability could be observed between different cells in a same coverslip, a balanced random selection of clusters across neurons, conditions and cultures was performed, and then variables from each culture were divided by the median of the control group. Results from different cultures were pooled and log(.) was applied, then the Mann-Whitney U value was computed. The process was repeated 1000 times and the p-value was determined from U distribution using the basic definition of the p-value. For SPT analysis, note that each QD is associated with 3 EA values, thus the sample size is 3 times greater. Statistical analysis were performed with R version 3.6.1 (R Core Team (2019). R: A language and environment for statistical computing. R Foundation for Statistical Computing, Vienna, Austria. Package used: ggplot2, matrixStats). Statistical tests were performed between a condition and its control only using cultures where both conditions were tested. Differences were considered significant for p-values less than 5% (*p < 0.05; **p < 0.01; ***p < 0.001; NS, not significant).

